# Bivalent binding of a fully human IgG to the SARS-CoV-2 spike proteins reveals mechanisms of potent neutralization

**DOI:** 10.1101/2020.07.14.203414

**Authors:** Bei Wang, Daniel Asarnow, Wen-Hsin Lee, Ching-Wen Huang, Bryan Faust, Patricia Miang Lon Ng, Eve Zi Xian Ngoh, Markus Bohn, David Bulkley, Andrés Pizzorno, Hwee Ching Tan, Chia Yin Lee, Rabiatul Adawiyah Minhat, Olivier Terrier, Mun Kuen Soh, Frannie Jiuyi Teo, Yvonne Yee Chin Yeap, Yuanyu Hu, Shirley Gek Kheng Seah, Sebastian Maurer-Stroh, Laurent Renia, Brendon John Hanson, Manuel Rosa-Calatrava, Aashish Manglik, Yifan Cheng, Charles S. Craik, Cheng-I Wang

**Affiliations:** Singapore Immunology Network, Agency for Science, Technology and Research (A*STAR), 8A Biomedical Grove, #03-06 Immunos, Singapore 138648, Singapore; Department of Biochemistry and Biophysics, University of California San Francisco (UCSF) School of Medicine, San Francisco, CA, USA; QBI COVID-19 Research Group (QCRG), San Francisco, CA, USA; Department of Pharmaceutical Chemistry, University of California San Francisco (UCSF), San Francisco, CA, USA; Virologie et Pathologie Humaine - VirPath team, Centre International de Recherche en Infectiologie (CIRI), INSERM U1111, CNRS UMR5308, ENS Lyon, Université Claude Bernard Lyon 1, Université de Lyon, Lyon, France; Biological Defence Program, DSO National Laboratories, 27 Medical Drive, Singapore 117510, Singapore; Bioinformatics Institute, Agency for Science, Technology and Research (A*STAR), 30 Biopolis Street, #07-01 Matrix, Singapore 138671, Singapore; School of Biological Sciences, Nanyang Technological University, Singapore, Singapore; Department of Microbiology and Immunology, Yong Loo Lin School of Medicine, National University of Singapore, Singapore, Singapore; VirNext, Faculté de Médecine RTH Laennec, Université Claude Bernard Lyon 1, Université de Lyon, Lyon, France; Department of Anesthesia and Perioperative Care, UCSF, San Francisco, CA, USA; Howard Hughes Medical Institute, UCSF, San Francisco, CA, USA

**Keywords:** SARS-CoV-2, COVID-19, SARS-CoV, coronavirus, spike protein, monoclonal antibody, receptor binding domain (RBD), phage display

## Abstract

*In vitro* antibody selection against pathogens from naïve combinatorial libraries can yield various classes of antigen-specific binders that are distinct from those evolved from natural infection^1–4^. Also, rapid neutralizing antibody discovery can be made possible by a strategy that selects for those interfering with pathogen and host interaction^5^. Here we report the discovery of antibodies that neutralize SARS-CoV-2, the virus responsible for the COVID-19 pandemic, from a highly diverse naïve human Fab library. Lead antibody 5A6 blocks the receptor binding domain (RBD) of the viral spike from binding to the host receptor angiotensin converting enzyme 2 (ACE2), neutralizes SARS-CoV-2 infection of Vero E6 cells, and reduces viral replication in reconstituted human nasal and bronchial epithelium models. 5A6 has a high occupancy on the viral surface and exerts its neutralization activity via a bivalent binding mode to the tip of two neighbouring RBDs at the ACE2 interaction interface, one in the “up” and the other in the “down” position, explaining its superior neutralization capacity. Furthermore, 5A6 is insensitive to several spike mutations identified in clinical isolates, including the D614G mutant that has become dominant worldwide. Our results suggest that 5A6 could be an effective prophylactic and therapeutic treatment of COVID-19.

---

The ongoing COVID-19 pandemic, caused by a novel coronavirus SARS-CoV-2 originating in Wuhan, China, at the end of 2019, is one of the world’s most devastating pandemics in living memory. As of July 1 2020, more than 10.7 million individuals have been infected worldwide resulting in nearly 518,000 deaths, and the numbers are still increasing. Currently only a single FDA-approved drug, Remdesivir, is available against COVID-19 and yet, it only shortened the duration of symptoms by 31% and had marginal improvement in survival, especially in the severely ill cases^6^. In this context, additional and more effective prophylactic and therapeutic drugs are urgently needed.

The coronavirus spike polypeptide comprises two subunits, S1 and S2, and is responsible for mediating viral entry^7^. Upon binding to the host cell receptor through its S1 subunit, the spike protein undergoes dramatic conformational changes and proteolytic processing, triggering the S2 subunit to facilitate fusion of viral and cellular membranes. Both SARS-CoV-2 and SARS-CoV use the angiotensin converting enzyme 2 (ACE2) as the entry receptor to infect host cells^8–10^. The receptor binding domain (RBD) located at the tip of the S1 subunit directly contacts ACE2 via the receptor binding motif (RBM), a small patch made up of about 20 amino acids^11, 12^ Thus, a rational approach to the development of the anti-SARS-COV or anti-SARS-COV-2 therapy is to explicitly direct antibody discovery against the RBD/ACE2 interaction interface^13^.

Since the first report of COVID-19 outbreak, discovery of SARS-CoV-2 neutralizing monocloncal antibodies (NAbs) has undergone an unprecedent speed. Some existing potent SARS-CoV NAbs that target the RBD/ACE2 interaction were quickly tested for their crossreactivity against SARS-CoV-2 because of the high similarity between the two viruses^10, 14, 15^, yet only very few showed potent neutralization^15–18^. Meanwhile, many SARS-CoV-2-specific NAbs have been discovered from immunized animals^19–21^ and COVID-19 convalescent patients^21–30^. Structural and biochemical studies indicated that most of these NAbs recognized epitopes within the ACE2 binding site, imposing direct competition between virus-ACE2 interaction^21, 22, 24, 25, 27, 28^. Other antibodies were found to bind to epitopes that are outside the RBM^18, 19, 21^ or even the RBD^23, 24^, suggesting that various mechanisms of neutralization exist. A number of these neutralizing antibodies have just entered clinical trials^21^.

In addition to those derived from natural infection or immunization, potent neutralizing antibodies against pathogens can be isolated by in vitro selection from highly diverse combinatorial human libraries^1–3^. Interestingly, some of such antibodies often came from different antibody subtypes and recognized different epitopes^31^. Following the release of the first SARS-CoV-2 genome, we set out to discover SARS CoV-2 neutralizing antibodies from a highly diverse human combinatorial antbody libray using a mechanism-based screening strategy. Here, we report the discovery of a potent NAb 5A6 with functional characterization in CHO-ACE2, Vero E6 cell line, and reconstituted human respiratory epithelium models, and reveal its molecular mechanism of action through cryogenic electron microscopy (cryo-EM). Furthermore, the cross-neutralizing activity of 5A6 against major SARS-CoV-2 mutants is demonstrated, further highlighting its potential as a COVID-19 therapy.

## Results and Discussions

### Isolation and characterization of SARS-CoV-2 neutralizing Fabs from a naïve human Fab phage display library

The naïve human Fab phage display library used in this project was constructed using the B cells donated by 22 healthy individuals, and has a diversity of 3 × 10^10^ clones^32^. To isolate neutralizing antibodies, we directed antibody selection against the recombinant SARS-CoV-2 S protein RBD^14^, the domain responsible for viral attachment to the host receptor ACE2 as previously discussed. Following stringent *in vitro* selection conditions and only two rounds of biopanning, ELISA screening of 570 individual clones identified 243 Fabs that were able to bind to the SARS-CoV-2 spike RBD. While several anti-SARS-CoV-2 antibodies have been reported to neutralize the virus by unclear mechanisms^19, 26^, to be expeditious, we took a direct, mechanism-based approach by screening for those Fabs that block the RBD/ACE2 interaction in a competition ELISA. Fabs were incubated with RBD before transfer to ELISA plate wells coated with the recombinant ACE2 protein, and the extent of RBD/ACE2 blocking was determined by the presence of ACE2-bound RBD. Twenty-seven Fabs with unique sequences exhibiting various degrees of blocking were identified (**Fig. 1a**). Binding characteristics of these Fabs to the SARS-CoV-2 spike RBD protein are shown in **Figure 1b**. Of the 27 selected clones, 19 clones were reformatted into full length IgGs, expressed recombinantly in CHO cells, and their binding to the RBD confirmed by concentration-dependent ELISA (**Extended Data Fig. 1**). Six antibodies, clones 1F4, 2H4, 3D11, 3F11, 5A6 and 6F8, showed strong receptor blocking by bio-layer interferometry (BLI) (**Fig. 1c**) and in competition ELISA, with IC_50_ values ranging between 0.4 nM and 36.8 nM (**Extended Data Fig. 2**). All six completely blocked the RBD/ACE2 interaction at high concentrations, except clone 3D11 which plateaued at 50%. Two Fabs, 5A6 and 3D11 showed the highest affinities for RBD at 7.6 nM and 1.6 nM respectively, presumably because of their slower relative off rates (**Fig. 1d, Extended Data Fig. 3**). The Fabs had intrinsic affinity varying from 1.6 nM (3D11) to 120 nM (3F11). They also varied by two orders of magnitude in their association and dissociation rates with 3F11 having the fastest on-rate and 3D11 having the slowest off-rate. The IgG format showed strong, subnano- or even picomolar binding avidity to the SARS-CoV-2 spike RBD either by BLI (**Fig. 1e**, **Extended Data Fig. 4**) or by ELISA (**Extended Data Fig. 5**). Interestingly, only 3D11 was able to cross react with SARS-CoV spike RBD, although the binding was much weaker (**Extended Data Fig. 5**). The data suggest that 3D11 recognizes a distinct epitope that is different from the other five antibodies and is conserved between SARS-CoV-2 and SARS-CoV. Indeed, stepwise binding BLI assays confirmed that 5A6 and 3D11 have non-overlapping footprints on the RBD while 5A6 shares overlapping epitopes with the other four antibodies (**Fig. 1f**).

**Fig. 1.**
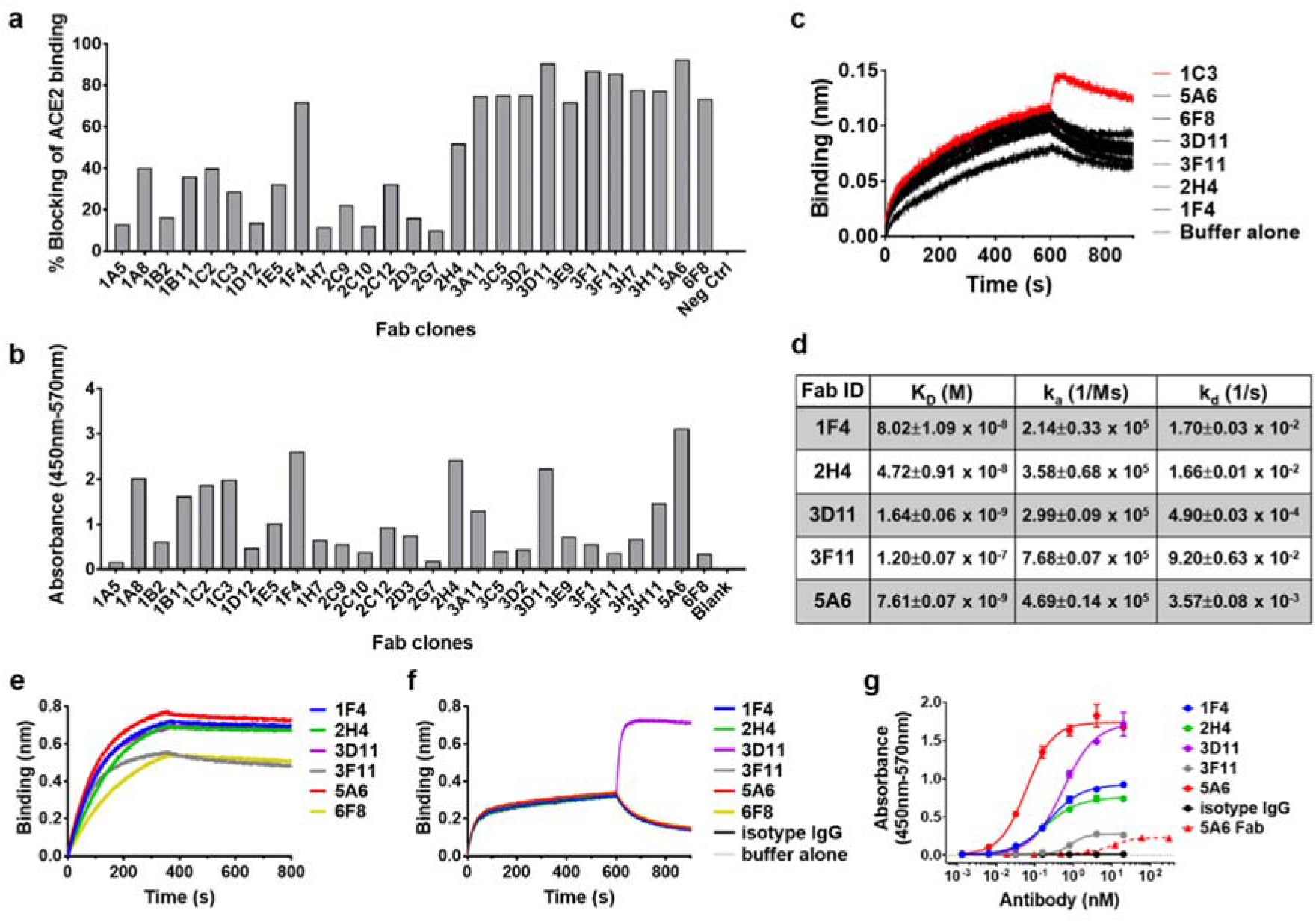
Identification of anti-SARS-CoV-2 spike RBD antibodies that can block ACE2/RBD interactions. **a**, Blocking of ACE2/RBD (SARS-CoV-2) interactions by 27 Fab clones, tested by competition ELISA. **b**, Binding avidity of the 27 Fab clones to SARS-CoV-2 spike RBD protein by ELISA. **c**, Competition of RBD-binding between ACE2 and six IgGs by biolayer interferometry assay. A weak blocking IgG clone 1C3 was included as a negative control. **d**, Binding affinity and the rate of association and dissociation of 5 Fab clones to SARS-CoV-2 spike RBD protein by BLI. Values are the means ± SD of two independent experiments. e, Binding avidity of six IgGs to the RBD by biolayer interferometry. The IgG concentration at 12.5 nM is shown here. **f**, Epitope binning of 5A6 by BLI analysis. Buffer alone, an isotype IgG and 5A6 IgG were included as controls. **g**, Binding of the IgG (solid lines, circles) and 5A6 Fab (dashed line, red triangle) to the purified SARS-CoV-2 pseudovirus. Data are presented as means ± SD in duplicates and are representative of two independent experiments.

To evaluate the neutralization potential of these SARS-CoV-2 RBD reactive antibodies, we generated pseudovirus expressing SARS-CoV-2 or SARS-CoV spike glycoprotein tagged with a luciferase reporter, and established a pseudovirus neutralization assay featuring CHO cells stably expressing human ACE2 (CHO-ACE2) as viral targets. All six antibodies showed dose-dependent neutralization (**Fig. 2a**). 5A6 strongly neutralized the SARS-CoV-2 pseudovirus with an IC_50_ of 75.5 ng/ml, followed by 1F4 (108 ng/ml), 2H4 (109 ng/ml), 6F8 (428.3 ng/ml), and 3F11 (930.4 ng/ml). Interestingly, while 3D11 neutralized SARS-CoV-2 pseudovirus, the neutralization did not follow the typical sigmoidal dose response, suggesting a unique mechanism of inhibition. Consistent with our binding studies, none of these antibodies showed significant neutralization of the SARS-CoV pseudovirus, suggesting high specificity of our antibodies to SARS-CoV-2. Interestingly, although 3D11 was able to bind weakly to the SARS-CoV RBD (**Extended Data Fig. 5**), it was unable to significantly neutralize SARS-CoV pseudovirus (**Fig. 2a**). These six antibodies were further evaluated in Vero E6 cells for neutralizing activity against a clinical SARS-CoV-2 strain, isolated from a patient in Singapore^33^. The 5 antibodies other than 3F11 showed >90% neutralization at the highest concentration (12.5 μg/ml), with 5A6 exhibiting an IC_50_ of 140.7 ng/ml, indicating >10-fold higher neutralizing potency than the other clones (**Fig. 2b**). In addition, 5A6 also potently neutralized a viral strain isolated in France with an IC_50_ of approximately 5 ng/ml (**Fig. 2c**). The difference between patient isolates is of unclear origin as they share identical spike sequences, but may be due to intrinsic properties such as rate of replication controlled by other viral genes.

**Fig. 2.**
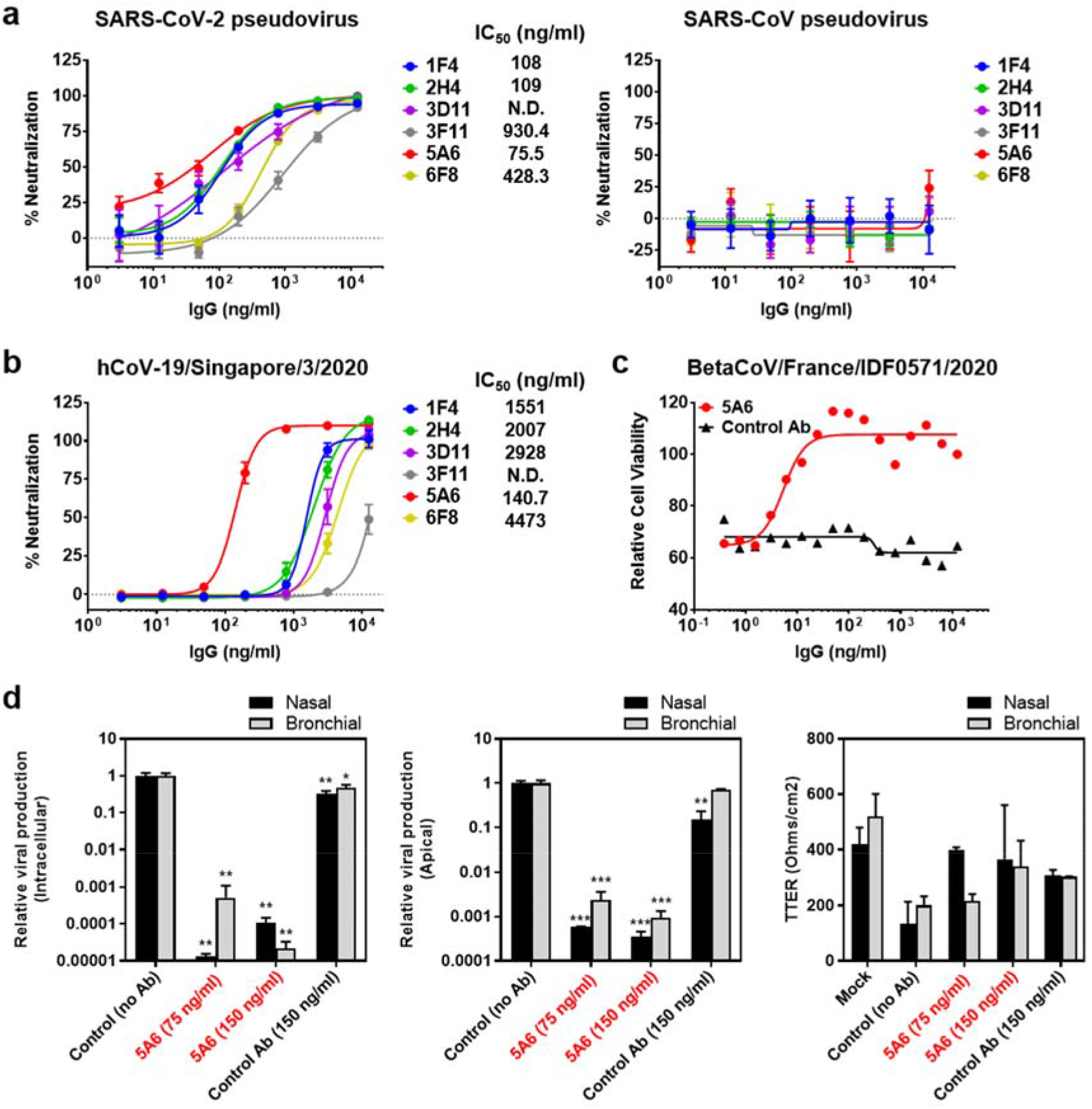
Neutralization of the SARS-CoV-2 pseudovirus and live viruses by anti-SARS-CoV-2 spike RBD IgG antibodies. **a**, Infection of CHO-ACE2 cells by SARS-CoV-2 pseudovirus (left panel) and SARS-CoV pseudovirus (right panel) were determined in the presence of 1F4, 2H4, 3D11, 3F11, 5A6 and 6F8 recombinant IgGs. Luciferase activities in the CHO-ACE2 cells were measured, and the percent neutralization was calculated. Data are presented as means ± SEM in triplicates and are representative of two independent experiments. **b**, Infection of Vero E6 C1008 cells by SARS-CoV-2 live virus (hCoV-19/Singapore/3/2020) were determined in the presence of 1F4, 2H4, 3D11, 3F11, 5A6 and 6F8 recombinant IgG. Infection induced cytopathic effect was determined by detecting the amount of ATP present in the uninfected live cells from which the percent neutralization was calculated. Data are presented as means ± SEM in triplicates and are representative of two independent experiments. **c**, Infection of Vero E6 cells by SARS-CoV-2 live virus (BetaCoV/France/IDF0571/2020) were determined in the presence of 5A6 and an isotype control IgG. Infection induced cytopathic effect was determined by detecting viable uninfected cells using MTS assay from which the percent neutralization was calculated. **d**, Evaluation of antiviral activity of 5A6 in a model of reconstituted human airway epithelia (HAE). Viral genome quantification was performed using RT-qPCR, and results presenting relative viral production (intracellular or apical) are expressed compared to control. Bars represent the means ± SD in duplicates. *\*\*\*P <0.001, ** P<0.01 and *P <0.05* compared to the control (no Ab) by one-way ANOVA. The trans-epithelial resistance (TEER in Ohms/cm2) was measures at 48hpi.

To explore the therapeutic potential of 5A6, we studied its neutralizing potency in physiologically relevant *in vitro* models. Human airway epithelia reconstituted using donor-derived, primary differentiated nasal and bronchial cells (MucilAir™ HAE^34^) were infected with SARS-CoV-2 virus in the presence or absence of 5A6 or an equivalent isotype control IgG. Viral replication was quantified as the measured copy number of the viral genomes inside, and at the apical poles of, nasal and bronchial HAE. **Figure 2d** shows that viral replication was drastically reduced, being 10^3^ to 10^4^-fold lower in the presence of 5A6 (75 or 150 ng/ml), while control IgG had little-to-no effect on infection. In addition, 5A6 also helped maintain epithelium integrity in the presence of virus, as evidenced by measurements of trans-epithelia electrical resistance. These results provided strong evidence in favour of the therapeutic potential of 5A6.

### Mechanism of SARS-CoV-2 neutralization by 5A6

Given that viral neutralization IC_50_s are more similar in general to the binding avidity than to the intrinsic affinity, we hypothesized that 5A6 neutralized the virus via bivalent binding to spike proteins on the viral surface. To support this hypothesis, we compared the neutralization potencies of the monovalent Fab to the bivalent IgG. A pseudovirus neutralization assay showed that the monovalent 5A6 Fab (IC_50_=663.9 ng/m, **Extended Data Fig. 6a**) had a 10-fold reduction in the neutralization potency relative to the bivalent 5A6 IgG (IC_50_=75.5 ng/ml, **Fig. 2a**). Reduction of neutralizing potency was also observed for the other Fabs (**Extended Data Fig. 6a**). Furthermore, the neutralization potency of the Fabs was also drastically reduced or even diminished in a live virus neutralization assay (**Extended Data Fig. 6b**), consistent with results using pseudovirus. This suggested that 5A6 IgG neutralizes the SARS-CoV-2 virus via bivalent binding to the spike protein or by crosslinking multiple spike protein trimers. However, while IgGs 1F4, 2H4, 3D11, 3F11 and 6F8 have similar or higher avidity for immobilized RBD, 5A6 has much higher neutralizing potency against both pseudovirus and live SARS-CoV-2 viruses. We asked if the discrepancy could arise from the differences in structural arrangement between the artificially immobilized RBD proteins and the natural RBD displayed on the viral surface. To answer this question, we purified the SARS-CoV-2 pseudovirus by gradient centrifugation and immobilized the viral particles on the ELISA plates. Concentration-dependent binding curves revealed that 5A6 IgG was packed on the viral surface at a much higher density than the other four IgG antibodies or 5A6 Fab, as shown by the higher optical signal (**Fig. 1g**). This may be a result of a unique binding mode that allows 5A6 IgG to accommodate a denser structural arrangement on the viral surface. To this end, it has been well documented that effective viral neutralization requires antibody packing density exceeding a critical threshold^35–37^, a criterion that was perhaps not easily achieved by the other five antibodies.

### Structural basis of viral neutralization by antibody 5A6

To better understand the molecular mechanism of 5A6 bioactivity, we determined single-particle cryo-EM structures of the trimeric spike protein alone at 2.4 Å resolution (not shown) and in complex with 5A6 Fab at 2.4 Å (**Fig. 3, Supplementary Movie 1, and Extended Data Fig. 7**). In the uncomplexed spike protein structure, individual RBDs are seen adopting two major conformations, here termed “up” and “down.” As previously reported, the resolution of an apo RBD in the up conformation is limited by its flexibility^8^. In the spike protein-5A6 Fab complex structure, the Fab hypervariable loops bind directly to surface loops at the tip of RBD. The RBD binding epitope of 5A6 (850 Å^2^) overlaps the binding site of the ACE2 receptor suggesting direct, competitive inhibition of ACE2 binding and confirming the mechanism based biopanning strategy for identification of ACE2 blockers (**Fig. 3b**). 3D classification of the spike protein-5A6 Fab complex dataset reveals several different subclasses in which the Fab occupancies in the trimer varies with two hallmark features. One striking highlight is that 5A6 Fab can bind to RBD in both up and down conformations while engaging the same epitope (**Fig. 3a-c**). While the orientations of the three RBDs within the trimer are different, the atomic resolution model built from one 5A6-RBD interface is superimposable with the other 5A6-RBD structures, and no steric clashes result from modelling any configuration of the complex. The resolution of our structure at the interface between 5A6 Fab and RBD is sufficiently high (~2.7 Å local resolution) to allow accurate modelling of the 5A6/RBD interface (**Fig. 3d**). Four of the six complementary determining regions (CDRs) of 5A6, namely, loops H1, H2, H3 and L1 are involved in binding to RBD. A surfeit of tyrosines from H1 (Tyr 32), H2 (Tyr 53 and Tyr 59), L1 (Tyr 32), and L3 (Tyr 92), as well as Arg 98, Thr 101, Val 103 and Arg 104 in the H3 loop cooperate to stabilize the interface. These interactions provide a structural framework that suggests why 5A6 is not affected by the V483I mutation while Leu 94 in the L3 loop is a potential liability in binding to the 483A mutation.

**Fig. 3.**
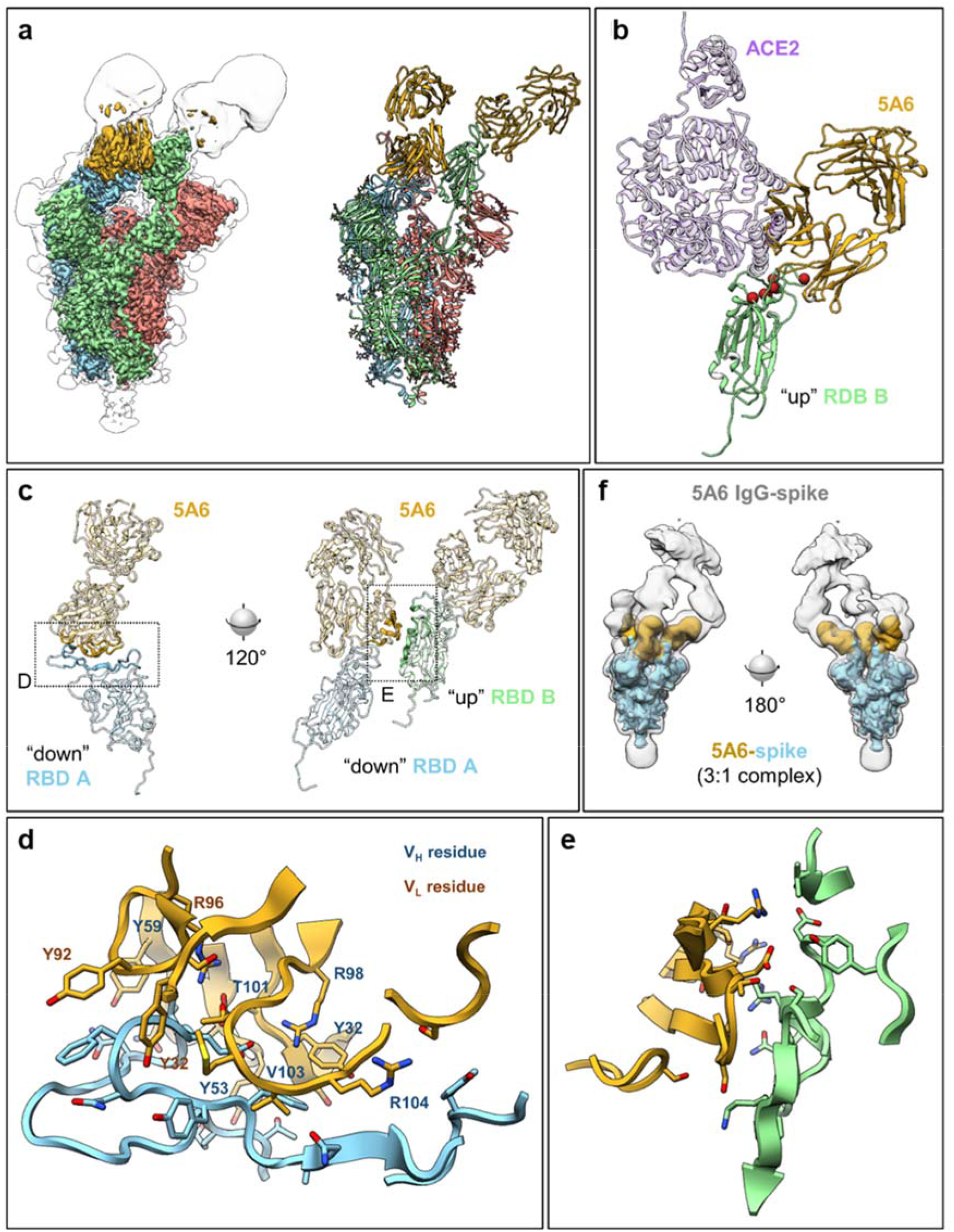
Cryo EM structures. **a**, Cryo-EM density (left) and model (right) of 5A6 bound to spike protein in a 2:1 complex (2.4 Å global resolution). Spike monomers A, B, & C are shown in blue, green and red, respectively, with two 5A6 Fabs in goldenrod (bound to A, “down,” and B, “up”). A second copy of the map is shown in transparent gray using a lower isovalue, to show the contour of the entire particle including flexible & low-resolution regions. See **Supplementary Movie 1** for additional viewing angles. **b**, Structural basis for virus neutralization by 5A6 binding to the ACE2-spike interface. Host receptor ACE2 (PDB 6m17, purple) is overlaid on the 5A6 (goldenrod) bound to spike RBD (green), revealing occlusion of the ACE2 binding site by 5A6. Red spheres indicate previously identified RBD mutations (N439, L455, F456, G476, V483, S494, and D614). Only V483 lies directly in the epitope. **c**, Fab 5A6 recognizes a complex tertiary epitope, and preferentially binds in a 2:1 complex to adjacent RBDs, one in the “down” conformation (spike monomer A) and the other in the “up” conformation (monomer B). The third RBD may be in either conformation, and either free or bound to another Fab. The A monomer with Fab bound to the epitope is shown left. A 120° rotation around the C3 axis, to the B monomer, reveals a secondary contact between the Fab light chain and B monomer (right). Dotted boxes show the regions focused in panels E and E, as indicated, and colors match those previous panels. **d**, Close-up of the 5A6 epitope corresponding to the labelled box in panel C. Fab residues mentioned in the text are indicated with dark blue labels for heavy chain, and dark orange for light chain. **e**, Close-up of the secondary contact between 5A6 light chain and the adjacent RBD, corresponding to the labelled box in panel C. Several close contacts and hydrogen bonds are formed at this additional interface, likely responsible for stabilizing the conformation of spike monomer A (“down” RBD). **f**, Cryo-EM density of an IgG bearing two copies of 5A6 bound to the spike trimer at ~15 Å resolution, with a binding mode congruent to that shown for the 5A6 Fab in the other panels.

The second hallmark feature is that two 5A6 Fabs are always found binding two neighbouring RBDs, one in an up position and one in a down position. A region of the light chain variable sequence of 5A6 that binds to the RBD in the down position forms a second interaction interface of 363 Å^2^ with a neighbouring RBD in the up position (**Fig. 3e**). This interaction presumably contributes to locking the first RBD in the down conformation, and further stabilizes RBD in the up conformation, as it is better resolved than other RBDs in the up position with or without Fab bound. Notably, the constant domain of 5A6 bound to the down RBD unavoidably clashes with the model of ACE2 bound to the neighbouring up RBD (**Fig. 3b** left), perhaps further potentiating neutralization.

The distances between all Fabs bound to a single trimer are sufficiently close to allow two Fabs from a single IgG to bind simultaneously to two different RBDs within the trimer, and we reasoned that binding mode might explain the strong avidity and high surface density of 5A6 IgG discussed above. We therefore applied single-particle cryo-EM to the spike protein complexed with 5A6 IgG, and the resulting ~15 Å structure reveals that 5A6 IgG indeed binds to two RBDs from the same trimer (**Fig. 3f**). While the Fc domain of the IgG is not well resolved, due to conformational flexibility and the tendency of the IgG to induce spike aggregation, the density clearly indicates that two Fabs from the same IgG prefer binding to two neighbouring RBDs in up and down conformations, just as described above for free 5A6 Fab. Thus, one spike protein trimer can accommodate 1.5 IgGs with a single IgG binding two RBDs and the third RBD bound to another IgG, potentially bridging separate spike protein trimers.

Remarkably, 5A6 directly competes with ACE2, binds any configuration of trimer RBD conformations without steric hindrance from either the spike protein or other Fabs, permits binding of IgG with unaltered geometry, and may disrupt cooperative opening of the entire trimer by locking adjacent RBDs into up and down conformations.

### Sensitivity of 5A6 and 3D11 to RBD mutations

A number of mutations resulting in amino acid changes in positions at or near the ACE2 interface have been isolated within more than 40,000 SARS-CoV-2 genomes sequenced to date. These include N439K in 246 samples (244 Scottish, 1 England, 1 Romania), V483A in 30 samples (26 USA/WA, 2 USA/UN, 1 USA/CT, 1 England), G476S in 18 samples (13 USA/WA, 2 USA/OR, 1 USA/ID, 1 USA/CT, 1 Belgium), S494P in 7 samples (3 USA/MI, 1 England, 1 Spain, 1 India, 1 Sweden), V483I in 2 English samples, and L455I together with F456V in one Brazilian sample. To determine if any of these mutant viruses could escape neutralization by 5A6 or 3D11, we produced the recombinant mutant RBD proteins to determine changes in binding avidity, and constructed mutant SARS-CoV-2 pseudoviruses to evaluate changes in neutralization potency. 5A6 IgG had a 4-fold reduction in binding avidity to the V483A mutation but was insensitive to the other five mutations (**Fig. 4a, Extended Data Fig. 8a**). Consistent with the binding results, 5A6 significantly lost neutralizing capacity against the V483A mutant pseudovirus. However, 5A6 largely retains activity against the other variants (**Fig. 4c**), including the D614G mutation that has spread at an alarming rate and become the dominant pandemic strain with a global frequency of 70.5% (GISAID), since its first identification in Europe in March^38^. Interestingly, 3D11 binding was not affected by any of these mutations (**Fig. 4b, Extended Data Fig. 8b**), but its neutralizing potency against the D614G mutant was partially reduced (**Fig. 4d**). The difference between the activity profiles of 5A6 and 3D11 could be explained by the different epitopes recognized by each antibody (**Fig. 1f**), and may argue for an antibody combination therapy to widen the neutralization spectrum against various mutant strains. While 3D11 has a weaker neutralizing potency than 5A6 in general, it can be further improved through antibody engineering, particularly via structure-guided approaches once high-resolution structures of its spike protein complex are determined. Hence, 3D11 could potentially rescue the loss of 5A6 neutralization potency in a virus strain bearing the V483A mutation, if these two antibodies are used in combination. Further structure-aided engineering is underway to broaden the neutralizing spectrum of 5A6 against other mutations.

**Fig. 4.**
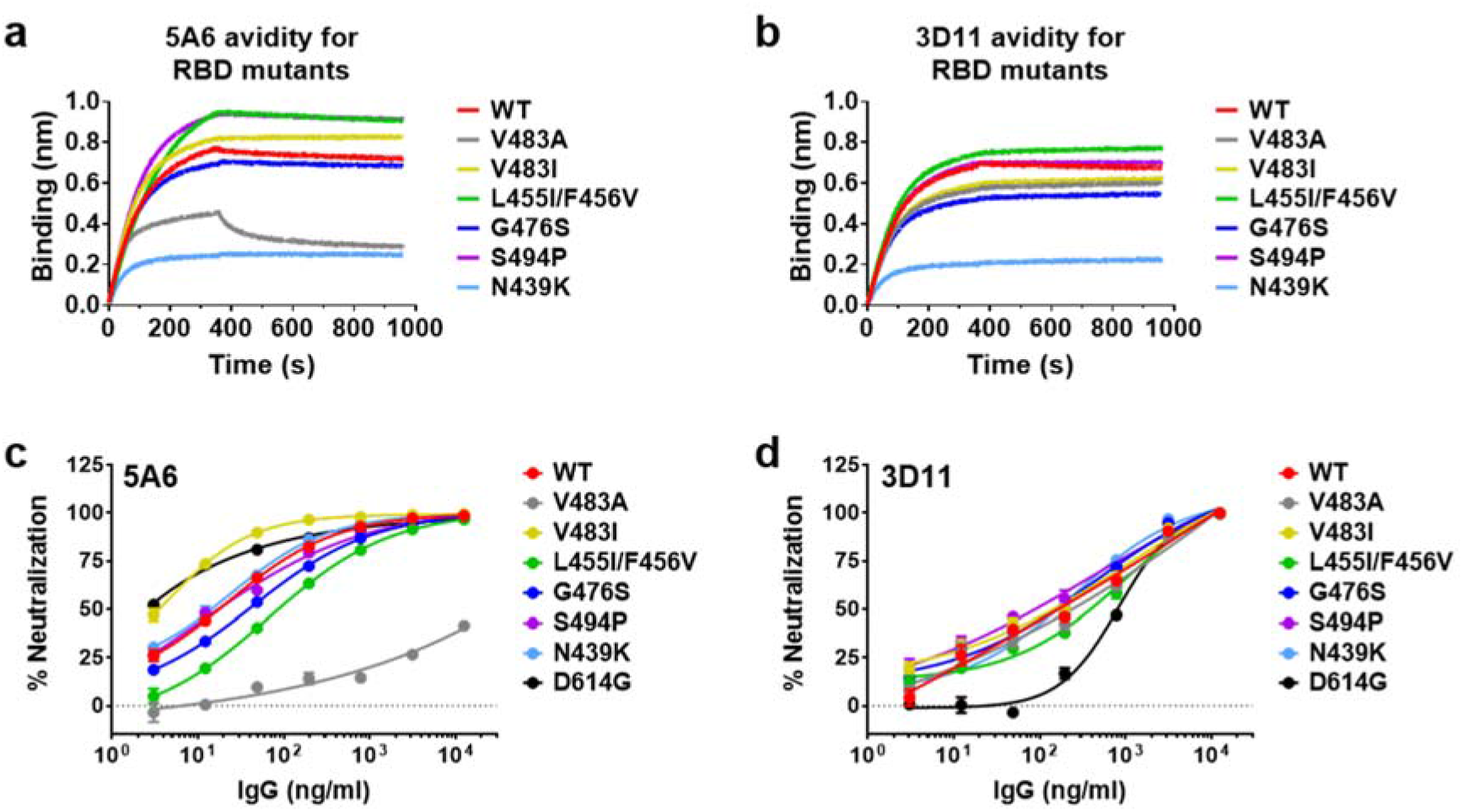
Sensitivity of 5A6 and 3D11 IgGs to RBD mutations. **a**, Binding avidity of IgGs 5A6 and **b**, to the wildtype RBD and six RBD mutants was determined by biolayer interferometry. IgG concentration at 12.5 nM is shown here. Neutralization of the SARS-CoV-2 pseudovirus with either wildtype (WT) or RBD mutations by **c**, 5A6 and **d**, 3D11 IgGs. Data are presented as means ± SEM in triplicates and are representative of two independent experiments.

In summary, our study demonstrates that potent anti-SARS-CoV-2 neutralizing antibodies can be rapidly discovered within a highly diverse, naïve combinatorial human antibody library using a mechanism-based screening strategy. Antibody 5A6 neutralizes SARS-CoV-2 by simultaneously binding to two RBDs of different structural orientations in the spike trimer, and prevents ACE2 binding by direct competition. The high resolution structures of the 5A6/RBD complex reveal details of extensive tertiary interactions, and greater neutralization potency, including against viruses with variant RBDs, can likely be achieved through structure-guided engineering, paving the way for the development of effective therapeutics for COVID-19.

## Acknowledgements

We thank Professor Yee-Joo Tan (Department of Microbiology, NUS; Institute of Molecular and Cell Biology, A*STAR) who kindly provided CHO-ACE2 cells and pXJ3’-S plasmid. We also thank Wong Pui San and Chye De Ho in the BSL 3 facility (DSO National Laboratories), as well as Dr. Pei Yin Lim (Singapore Immunology Network) for their technical assistance. We thank Professor Jason McLellan (University of Texas at Austin) for the prefusion S ectodomain. We thank Professor Joseph DeRisi and Professor Robert Fletterick (UCSF) and Dr. John Pak (CZI Biohub, San Francisco) for helpful discussions. This work was supported by core research grants provided to Singapore Immunology Network of the Biomedical Research Council (BMRC), the A*CCLERATE GAP fund (ACCL/19-GAP064-R20H-I) from the Agency of Science, Technology and Research (A*STAR, Singapore). Support was also provided by the National Institutes of Health Grant P50AI150476 (CSC, YC, MB), NIH Grants S10 S10OD020054 (YC) and S10OD021741 (YC) and the UCSF Program for Breakthrough Biomedical Research which is partially funded by the Sandler Foundation (C.S.C., Y.C., A. M.). Y.C. is an Investigator of the Howard Hughes Medical Institute.

## Author contributions

B.W., D.A., W.H.L., C.W.H., B.F., P.M.L.N., E.Z.X.N., M.B., D.B., S.M.S., M.R.C., A.M., Y.C., C.S.C., and C.I.W. conceptualized and designed the study. L.R., B. J.H., M.R.C., A.M., Y.C., C.S.C., and C.I.W. coordinated the study. B.W., D.A., W.H.L., C. W.H., B.F., P.M.L.N., E.Z.X.N., M.B., D.B., H.C.T., C.Y.L., R.A.M., A.P., O.T., M.K.S., F.J.T., Y.Y.C.Y., Y.H., and S.G.K.S. conducted the experiments. B.W., D.A., W.H.L., C.W.H., B. F., P.M.L.N., E.Z.X.N., M.B., D.B., H.C.T., C.Y.L., R.A.M., A.P., O.T., S.M.S, L.R., M.R.C., A.M., Y.C., C.S.C., and C.I.W. analysed and interpreted the data. B.W., D.A., W.H.L., xC. W.H., B.F., P.M.L.N., M.B., D.B., A.P., O.T., S.M.S., L.R., B.J.H., M.R.C., A.M., Y.C., C.S.C., and C.I.W. wrote and revised the manuscript.

## Competing interests

B.W., W.H.L., C.W.H., P.M.L.N., E.Z.X.N., H.C.T., C.Y.L., R.A.M., M.K.S., F.J.T., Y.Y.C.Y., Y.H., and C.I.W. are listed as inventors of a filed patent for all 27 monoclonal antibodies mentioned in this manuscript. The other authors declare that they have no competing interests.

## Data and materials availability

All data are available in the manuscript or in the supplementary materials and the source data are provided as a Source Data file. All antibodies are proprietary and can be obtained through an Materials Transfer Agreement.

