## Supplementary Materials for "Bivalent binding of a fully human IgG to the SARS-CoV-2 spike proteins reveals mechanisms of potent neutralization"

**This file includes:**
Methods
Extended Data Fig.1 to 8
Extended Data Table 1
Legend to Suppelementary Movie 1

#### Methods

##### Antibody discovery from phage display library

Anti-SARS-CoV-2 spike RBD antibodies were isolated from an HX02 human Fab phage display library (Humanyx Pte Ltd) via *in vitro* selection. Briefly, biopanning was performed using SARS-CoV-2 RBD (YP_009724390.1) (Arg319-Phe541) with a mouse Fc tag (Sino Biological, 40592-V05H) biotinylated using the EZ-Link NHS-PEG4-Biotin labelling kit (Thermo Fisher Scientific, #A39259). In both rounds of biopanning, biotinylated SARS-CoV-2 RBD-mFc protein was immobilized on M280 streptavidin-coated magnetic beads (Life Technologies, #11205D); 3.5 x $10^{12}$ cfu phage in 1mL 1% casein-PBS blocking buffer was used in the first round, and 1.64 x $10^{11}$ cfu phage were used in the second round. During the biopanning process, binders to mouse Fc were removed by pre-incubation of phage with 2 µM mouse IgG before mixing with the RBD-mFc antigen. After two rounds of biopanning, the Fabs of selected clones were expressed in *E. coli* HB2151 cells (Stratagene) to screen for RBD binders by ELISA. Unique clones were identified by DNA sequencing.

##### IgG expression and purification

Fabs were reformatted into human IgG in the pTT5 vector (National Research Council of Canada) and the IgG antibodies were expressed using ExpiCHO expression system (Thermo Fisher Scientific) by transient co-transfection of plasmids expressing the heavy and light chain of each antibody clone. Eight days after transfection, ExpiCHO-S cell suspension was centrifuged for 10 min at 2000 rpm and filtered with 0.22 µm filter to remove the cells and debris. Antibodies were then purified from the culture supernatant using Protein G resin (Merck Millipore) following the manufacturer's instructions. After elution, the purified antibodies were dialyzed at 4°C for 4-20 hours against 1x PBS, for 3 times and concentrated to 1-2 mg/ml using 10MWCO Vivaspin 20 (Sartorius).

**Fab production and purification**

The tag-less Fab fragments were produced using the ExpiCHO transient expression system. Eight days after transfection, ExpiCHO-S cell suspension was centrifuged and filtered; and Fab was purified from the filtered culture supernatant using cation exchange chromatography (CIEX) on AKTA FPLC System (GE Healthcare). In brief, the supernatant was concentrated to 2 ml using 10MWCO Vivaspin 20 (Sartorius), diluted 1:20 in Buffer A (20 mM Sodium Acetate, pH 5.2), filtered through 0.22 µm filter, and loaded onto Mono-S 5/50 GL column at a flow rate of 1ml/min. Fab fragments were eluted in Buffer B (20 mM Sodium Acetate, pH 5.2 with 1 M Sodium Chloride) with a sequential linear gradient of 0% to 5% in 5 min, 5% to 15% in 30 min, and 15% to 100% in 20 min of Buffer B injection at a flow rate of 1 ml/min. The resulting purified Fab fragments were dialyzed at 4°C for 4-20 hours against 4 liters of 20 mM Histidine, 150 mM NaCl, pH 6.6, for 3 times and concentrated to 1-2 mg/ml using 10MWCO Vivaspin 6 (Sartorius).

**Avidity binding ELISA to recombinant RBD proteins**

Anti-SARS-CoV-2 spike RBD IgG antibodies were tested in an ELISA against biotinylated recombinant SARS-CoV-2 spike protein RBD-mFc (Sino Biological, 40592-V05H) or SARS-CoV spike protein RBD-His (Sino Biological, 40150-V08B2) to assess binding avidity for the target. In brief, NeutrAvidin protein (Thermo Fisher Scientific, #31000) was coated at 5 µg/ml onto 96-well ELISA plates in coating buffer (8.4 g/L $NaHCO_3$, 3.56 g/L $Na_2CO_3$, pH 9.5) overnight at 4°C. After blocking with 1% Casein (Thermo Fisher Scientific, #A37528) for two hours, biotinylated antigen at 0.2 µg/ml was added to the plates and captured by NeutrAvidin during one-hour incubation at room temperature. After washing with 0.05% PBST for 5 times, the IgG antibodies were added at different concentrations with 3-fold dilutions in triplicate and incubated for one hour. The wells were then washed again with 0.05% PBST, followed by addition of Peroxidase-conjugated AffiniPure F(ab')2 Fragment Goat Anti-Human IgG, Fcγ Fragment Specific (JACKSON ImmunoResearch, #109-036-098, 1:3000). Finally, the wells were washed and the HRP activity was measured at 450 nm with addition of 3,3',5,5'-tetramethylbenzidine (TMB) substrate (Surmodics, BioFX®, TMBW-1000-01).

**Avidity binding ELISA to purified pseudoviral particles**

Anti-SARS-CoV-2 spike RBD IgG antibodies were tested in an ELISA against iodixanol-gradient-purified SARS-CoV-2 psuedoviruses with an isotype IgG used as a negative control antibody. In brief, 1 μg/ml of psuedoviral particles were coated in coating buffer onto 96-well ELISA plates overnight at 4°C. After blocking with 1% Casein (Thermo Fisher Scientific, #A37528) for two hours, serially diluted IgG antibodies starting from 100 nM with five-fold dilutions were added to the plates and incubated for an hour at room temperature. The wells were then washed again with 0.05% PBST, followed by addition of HRP conjugated anti-human Fc antibody (Thermo Fisher Scientific, #A21445, 1:3000) for one-hour incubation before HRP activity was measured at 450 nm with addition of TMB substrate.

**Competition ELISA**

Anti-SARS-CoV-2 spike RBD IgG antibodies were tested in a competition ELISA to assess their ability to block the spike protein RBD from binding to human ACE2 protein. In brief, the recombinant human ACE2 protein with a human Fc tag (ACE2-Fc) was coated onto the 96-well ELISA plates in coating buffer overnight at 4°C and blocked with 1% Casein. Then different concentrations of anti-SARS-CoV-2 spike RBD IgG antibodies were pre-incubated with 0.5 nM biotinylated spike protein RBD-mFc for one hour at room temperature before they were added to the ELISA plates coated with ACE2-Fc. After one-hour incubation, the wells were washed with 0.05% PBST for five times and HRP conjugated streptavidin (Biolegend, #405210) was added at a dilution of 1:3000, and incubated for another one hour before HRP activity was measured at 450 nm with addition of TMB substrate.

**Cell lines and cell culture**

The human embryonic kidney epithelial cell 293T (ATCC, CRL-3216) was cultured in Dulbecco's modified Eagle's medium (Hyclone, SH30022.01) supplemented with 10% heat-inactivated FBS (Gibco, 10270-106). A stable cell line expressing human ACE2, CHO-ACE2 (a kind gift from Professor Yee-Joo Tan, IMCB, A*Star)[39] was maintained in Dulbecco's modified Eagle's medium supplemented with 10% heat-inactivated FBS, 1% MEM Non-Essential Amino Acids Solution (Gibco, 11140-050) and 0.5 mg/ml of Geneticin™ Selective Antibiotic (Gibco, 10131-027). Every 2-3 days, cells were passaged by dissociating the cells with StemPro™ Accutase™ Cell Dissociation Reagent (Gibco, A1110501).

**Conversion of IgG antibodies to Fab fragments**

Digestion reaction for each IgG was prepared using immobilized FabALACTICA microspin columns (Genovis). 100 μL of IgG at 5 mg/mL concentration in digestion buffer (150 mM sodium phosphate, pH 7.0) were incubated overnight on each column. Digested sample was further purified using a HiTrap Protein L column followed by size-exclusion chromatography (Superdex 75 10/300 GL) using an Äkta Pure FPLC (all GE Healthcare).

**Soluble SARS-CoV-2 Spike Production and Purification**

The expression plasmid containing the prefusion S ectodomain as used in Wrapp, et al.[40] was provided by the McLellan's lab, University of Texas at Austin. This construct was used to transiently transfect high-density Chinese Hamster Ovary (ExpiCHO) cells with

ExpiFectamine per the “Max Titer” protocol provided (Thermo Fisher). Six days post-transfection, 0.2 µm-filtered supernatant was collected and incubated with Ni-Sepharose Excel (Cytiva Life Sciences) for batch purification. Eluate was collected, concentrated in a 50 MWCO Amicon Ultra-15 centrifugal filter unit (MilliporeSigma), and injected onto a Superose6 10/300 GL column equilibrated in 10 mM HEPES, 200 mM NaCl, pH 8.0 to isolate trimeric, monodisperse material for Fab/IgG complexing.

**Sample preparation for cryo-EM**

2.5µL of Spike-Fab complex at a concentration of 0.4 mg/mL was applied to a 300 mesh gold Quantifoil 1.2/1.3 holey carbon grid that was glow discharged for 30 sec at 15 mA immediately before sample application. Grids were blotted using Whatman #1 filter paper for 8 or 10 seconds at a blot force of 0 at 4°C and 100% humidity using a Mark IV Vitrobot (Thermo Fisher) and plunge frozen into liquid ethane. Samples were loaded onto a Titan Krios transmission electron microscope (Thermo Fisher) equipped with a Gatan K3 direct electron detector (Gatan) and a Quantum GIF energy filter (Gatan) operated with a 20 eV slit width during image acquisition. The K3 camera was operated in CDS mode using super resolution. A nominal magnification of 105,000x was used, for a pixel size of 0.835 Å (0.4175 Å super resolution pixel size) at the sample. A dose rate of 8 $e^-/(pix \cdot sec)$, or 11.5 $e^-/(Å^2 \cdot sec)$, and a frame rate of 0.05 sec/frame was used with a total exposure time of 5.9 sec, for a total dose of 67.7 $e^-/Å^2$. Automated data collection was performed using SerialEM[41].

**Image processing**

Dose-weighted, motion-corrected sums down-sampled to the physical pixel size were obtained from the super-resolution DED movies using UCSF Motioncor2[42]. For the spike trimer, CTF estimation was performed in cryoSPARC[43] followed by blob-based particle picking, 2D classification, *ab initio* modelling, 3D classification, and 3D refinement. For images of antibody complexes, particles were instead picked using templates generated from the apo trimer structure, and the apo trimer was likewise used as an initial model in 3D classification. The resolution of the interface between the spike RBD and the 5A6 Fab was further improved using naïve focused refinements. Processing details are given in Extended Data Table 1 and Extended Data Figure 7.

**Molecular modelling**

The previously determined structure of the SARS-CoV-2 spike protein with one open RBD (PDB: 6vyb), along with a full-length homology model of 5A6 Fab computed by MODELLER[44], were simultaneously docked into cryo-EM density using UCSF Chimera[45]. Missing segments and side chains in the RBDs were built using Coot. Finally, real-space refinement in PHENIX[46], and density-restrained molecular dynamics simulations in ChimeraX[47] and ISOLDE[48] were used to finalize the model of the 5A6-RBD 2:1 complex. Model statistics and density fit are presented in **Extended Data** Table 1 and Extended Data Figure 7.

**Generation of pseudovirus particles expressing SARS-CoV-2 Spike glycoprotein**

Psuedotyped viral particles expressing SARS-CoV-2 or SARS-CoV spike protein were produced by transfecting of 30 million 293T cells with 12 µg pMDLg/pRRE (Addgene #12251), 6 µg pRSV-Rev (Addgene #12253), 24 µg pHIV-Luc-ZsGreen (Addgene #39196) and 12 µg pTT5LnX-coV-SP (expressing SARS-CoV-2 spike protein, Genbank: YP_009724390.1, a kind gift from DSO National Laboratories) or 12 µg pXJ3'-S (expressing SARS-CoV spike protein from HKU39849 strain, a kind gift from Professor Yee-Joo Tan, IMCB, A*STAR)[39] using Lipofectamine 2000 transfection reagent (Invitrogen, 11668-019).

The transfected cells were cultured at 37°C incubator for 3 days. Viral supernatant was harvested, centrifuged at 700 g for 10min to remove cell debris and filtered through a 0.45 µm filter unit (Sartorius #16555). Lenti-X p24 rapid titer kit (Takara Bio, #632200) was used to quantify the viral titres following the manufacturer instructions. Plasmids expressing SARS-CoV-2 spike proteins with 6 different RBD mutations (V483A, V483I, L455I/F456V, G476S, S494P, and N439K) as well as D614G mutation were generated using QuickChange Lightning Multi Site-Directed Mutagenesis Kit (Agilent, #210513) and the pseudovirus particles expressing these spike proteins with mutated RBD were produced.

**Concentration and purification of pseudovirus particles**

To concentrate and purify the pseudovirus particles expressing the SARS-CoV-2 spike glycoproteins with either wildtype RBD or V483A mutant RBD, pre-cleared 40 mL viral supernatant was concentrated by 20% sucrose gradient centrifugation at 10,000 g for 4 hours at 4°C in an SW41 Ti rotor with no brake. Upon removal of supernatant, 1mL of PBS was added to the virus pellet and left at 4°C overnight. Concentrated virus was further purified by an OptiPrep (60% [wt/vol] iodixanol; #07820; STEMCELL Technologies Inc) velocity gradient. Iodixanol gradients were prepared in PBS in 1.2% increments ranging from 6 to 18%. Psuedoviruses were layered onto the top of the gradient and centrifuged for 1.5 hours at 200,000 g in an SW41 Ti rotor. Gradient fraction that contained psuedovirus pellet was collected.

**Antibody neutralization assay with SARS-CoV-2 or SARS-CoV Spike glycoprotein pseudovirus**

CHO-ACE2 cells were seeded at a density of 3.2 x $10^4$ cells in 100 µL of complete medium without Geneticin in 96-well Flat Clear Bottom Black Polystyrene TC-treated Microplates

(Corning, #3904). Serially diluted IgGs were incubated in a 96-well flat-bottom cell culture plate (Costar, #3596) with an equal volume of pseudovirus (12 ng of p24) at the final volume of 50µL at 37°C for one hour, and the mixture was added to the monolayer of pre-seeded CHO-ACE2 cells in triplicate. After one hour of pseudovirus infection at 37°C, 150µl of culture medium was added to each well and the cells were further incubated for another 48 hours. Upon removal of culture medium, cells were washed twice with sterile PBS, and then lysed in 20 µL of 1x Passive lysis buffer (Promega, E1941) with gentle shaking at 400 rpm at 37°C for 30 minutes. Luciferase activity was then assessed using a Luciferase Assay System (Promega, E1510) on a Promega GloMax Luminometer. The relative luciferase units (RLU) were converted to percent neutralization and plotted with a non-linear regression curve fit using PRISM.

**Antibody neutralization assay with live SARS-CoV-2 virus in Vero E6 cells**

The potency of the six IgG antibodies, 1F4, 2H4, 3D11, 3F11 5A6 and 6F8 were determined in neutralizing live SARS-CoV-2 virus assays. In brief, 25 µl of 100 $TCID_{50}$ of SARS-CoV-2 live virus (hCoV-19/Singapore/3/2020) isolated from a nasopharyngeal swab of a patient in Singapore[33], was mixed with an equal volume of serially diluted IgG or Fab antibodies and incubated at 37°C for one hour before the mixture was added to 50 µl of Vero E6 C1008 cells in suspension. The infected cells were incubated at 37°C incubator for four days and the cell viability was determined using Viral ToxGlo™ Assay (Promega, #G8941). The potency of 5A6 IgG was also tested in neutralizing a live virus strain isolated from one of the first COVID-19 cases confirmed in France: a 47-year old female patient hospitalized in January 2020 in the Department of Infectious and Tropical Diseases, Bichat Claude Bernard Hospital, Paris[49]. The complete viral genome sequence was obtained using Illumina MiSeq sequencing technology, was then deposited after assembly on the GISAID EpiCoV platform (Accession ID

EPI_ISL_411218) under the name BetaCoV/France/IDF0571/2020. Similarly, 25 µl of 100 $TCID_{50}$ of SARS-CoV-2 live virus was mixed with an equal volume of serially diluted 5A6 IgG or control IgG and incubated at 37°C for one hour before the mixture was added to 50 µl of Vero E6 cells in suspension. The infected cells were incubated at 37°C incubator for four days and the cell viability was determined using the CellTiter 96® $AQ_{ueous}$ One Solution Cell Proliferation Assay (Promega, #G3582).

**Antibody neutralization assay with live SARS-CoV-2 virus in reconstituted human airway epithelia (HAE)**

MucilAir$^{TM}$ HAE reconstituted from human primary cells obtained from nasal or bronchial biopsies were provided by Epithelix SARL (Geneva, Switzerland) and maintained in air-liquid interphase with specific culture medium in Costar Transwell inserts (Corning, NY, USA) according to the manufacturer's instructions. For antibody neutralization assay, the apical poles of HAE were gently washed twice with warm Opti-MEM medium (Gibco, ThermoFisher Scientific) and then infected directly with a 150 µl dilution of live SARS-CoV-2 virus (strain: BetaCoV/France/IDF0571/2020) in Opti-MEM medium, at a multiplicity of infection (MOI) of 0.1. Viral suspensions were pre-incubated 60 min with anti-S antibody 5A6 IgG (75 ng/ml or 150 ng/ml) or an anti-Ebola glycoprotein control antibody (150 ng/ml) before infection. A control infection was performed in absence of antibody. For mock infection, the same procedure was followed using Opti-MEM as inoculum. Samples collected from apical washes at 48 hours post-infection were separated into 2 tubes: one for TCID50 viral titration (stored at -80°C) and one for RT-qPCR. HAE cells were harvested in RLT buffer (Qiagen) and total RNA was extracted using the RNeasy Mini Kit (Qiagen) for subsequent RT-qPCR. Variations in trans epithelial electrical resistance (Δ TEER) were measured using a dedicated volt-ohm meter (EVOM2, Epithelial Volt/Ohm Meter for TEER) and expressed as Ohm/cm$^2$.

**Fab affinity measurement by BioLayer Interferometry (BLI)**

Binding affinity of purified Fab to RBD was measured on the Octet96Red system (ForTeBio). Anti-human IgG Fc (AHC) sensors were first loaded with 1 μg/ml of Fc-RBD for 10 min, followed by kinetics buffer (phosphate-buffered saline buffer supplemented with 0.1% Tween-20 and 0.1% BSA) for 5 min to establish a stable baseline. The sensors were then dipped into different concentrations of each Fab from 100 nM to 3.125 nM in two-fold dilutions for 6 min, and then in kinetics buffer again for 10 min to measure association and dissociation. Assays were run at 25°C and data was analysed on the Octet System Data Acquisition Software version 9.0.0.4. using the 1:1 model.

**Avidity binding by BLI**

Avidity of anti-SARS-CoV-2 spike RBD IgG antibodies for RBD and various RBD mutants was measured on the Octet96Red system. Anti-hIgG Fc capture (AHC) sensors were used. The sensors were loaded with 1 μg/ml of Fc-RBD or Fc-RBD mutants (made in-house) in assay buffer (phosphate-buffered saline buffer supplemented with 0.1% Tween-20 and 0.1% BSA) for 10 min, quenched in 0.5 mg/ml of isotype IgG in assay buffer for 10 min, then dipped in assay buffer for 12 min for the system to stabilize. To measure the association of 5A6, the sensors were dipped in a range of 5A6 IgG concentrations (25-0.39 nM in 2-fold serial dilutions) in assay buffer for 6 min. To measure dissociation, the sensors were dipped in assay buffer for 10 min. The experiment was conducted at 25°C. Data analysis was done in the Octet System Data Acquisition Software version 9.0.0.4. using the 1:2 bivalent model.

**ACE2 competition assay by BLI**

Competition with ACE2 for RBD was measured on the Octet96Red system. The amine-reactive (AR2G) sensor tips (ForteBio) were activated in freshly prepared 20 mM EDC (1-ethyl-3-[3-dimethylaminopropyl]-carbodiimide hydrochloride), 10 mM NHS (N-hydroxysuccinimide) solution and ACE2-Fc was immobilized to the sensor tips using a concentration of 10 µg/ml in 10 mM sodium acetate pH4 buffer. After quenching in 1M ethanolamine, the ACE2-Fc-immobilized sensor tips were dipped in 5 µg/ml of tagless RBD for 600s then in 10µg/ml of the second antibody for 300s. The assay was run at 25°C.

**Epitope binning by BLI**

Epitope binning was done using a classical sandwich assay. The AR2G sensor tips (ForteBio) were activated in freshly prepared 20mM EDC (1-ethyl-3-[3-dimethylaminopropyl]-carbodiimide hydrochloride), 10mM NHS (N-hydroxysuccinimide) solution and the 5A6 antibody was immobilized to the sensor tips using a concentration of 7.5 µg/ml of 5A6 in 10 mM sodium acetate pH 6 buffer. After quenching in 1M ethanolamine, the 5A6-immobilized sensor tips were dipped in 5 µg/ml of tagless RBD for 600s, then in 10µg/ml of the second antibody for 300s. The assay was run at 25□C. Sensor tips were regenerated in 10 mM glycine at pH 2.7 and neutralized in PBS with 0.1% Tween-20 before another cycle of sandwich assay was performed. Each sensor tip was used in a total of 3 cycles. Data analysis was done in the Octet System Data Acquisition Software version 9.0.0.4.

**Statistical analysis.**

Data were analysed using GraphPad Prism version 7.03. Statistical tests are indicated in the figure legends. $EC_{50}$ values were calculated by non-linear regression analysis on the binding curves using GraphPad Prism and $IC_{50}$ values values were calculated using the [Inhibitor] vs response variable slope four parameter of GraphPad Prism. One-way analysis of variance

(ANOVA) was used to compare differences between groups. Differences were considered statistically significant at confidence levels $*P < 0.05$ or $**P < 0.01$, $***P < 0.001$.

### Extended data figures and table

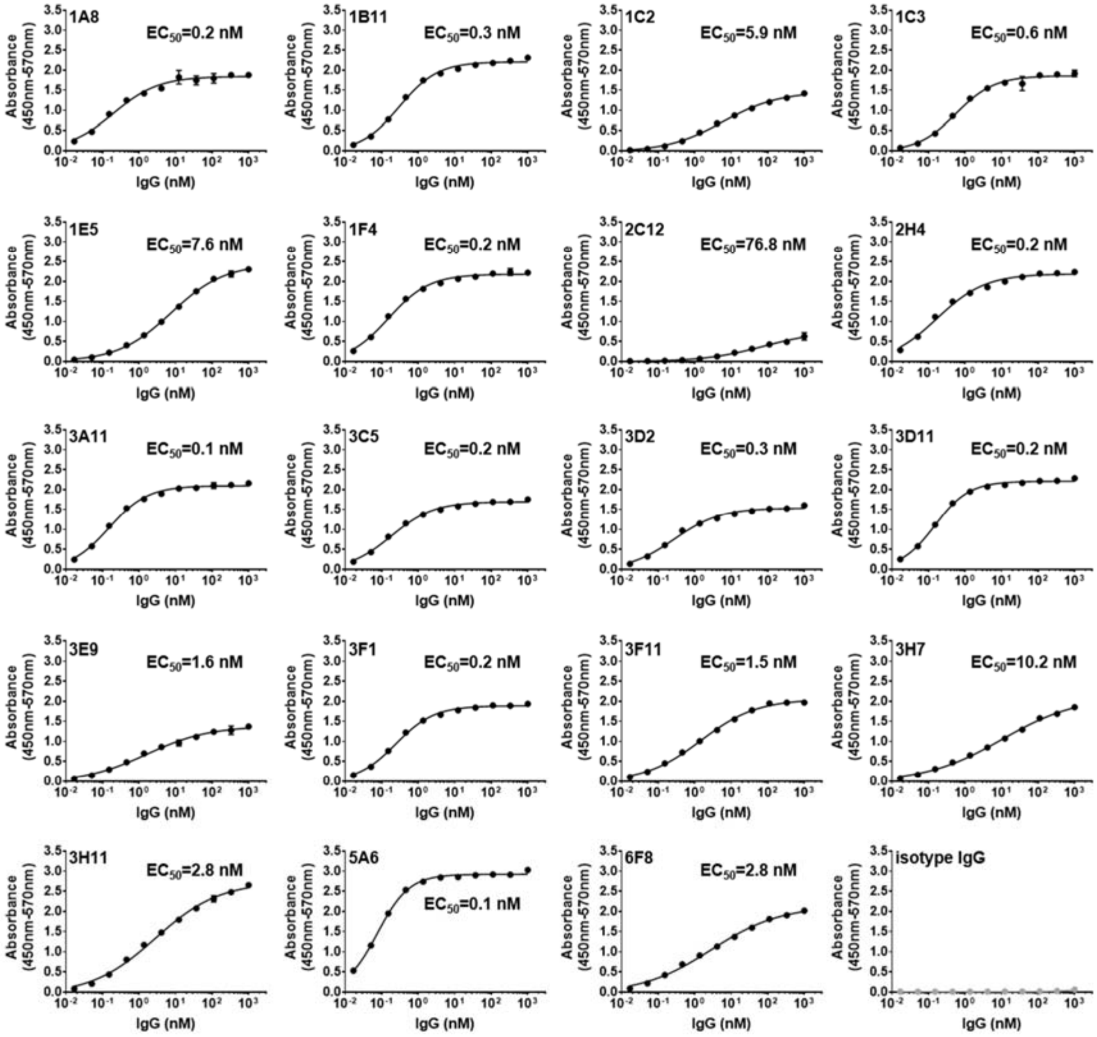

**Extended Data Fig. 1 Binding avidity of the 19 IgGs to SARS-CoV-2 spike RBD protein tested by ELISA.** Data are presented as means ± SD in duplicates.

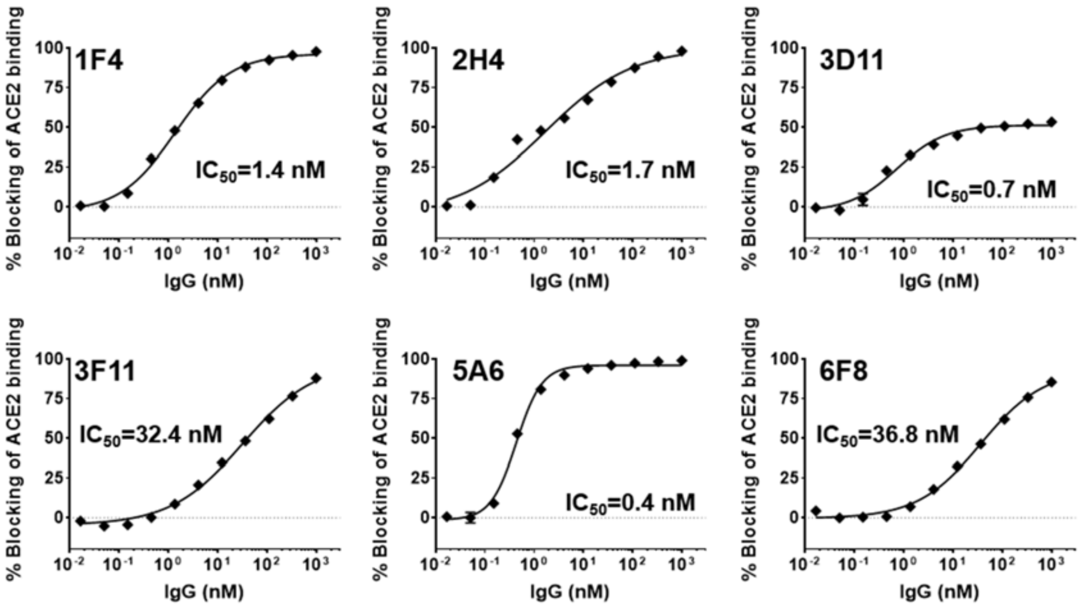

**Extended Data Fig. 2 Blocking of ACE2/ SARS-CoV-2 RBD interaction by 1F4, 2H4, 3D11, 3F11, 5A6 and 6F8 IgGs tested by competition ELISA.** Data are presented as means ± SD in triplicates and are representative of two independent experiments.

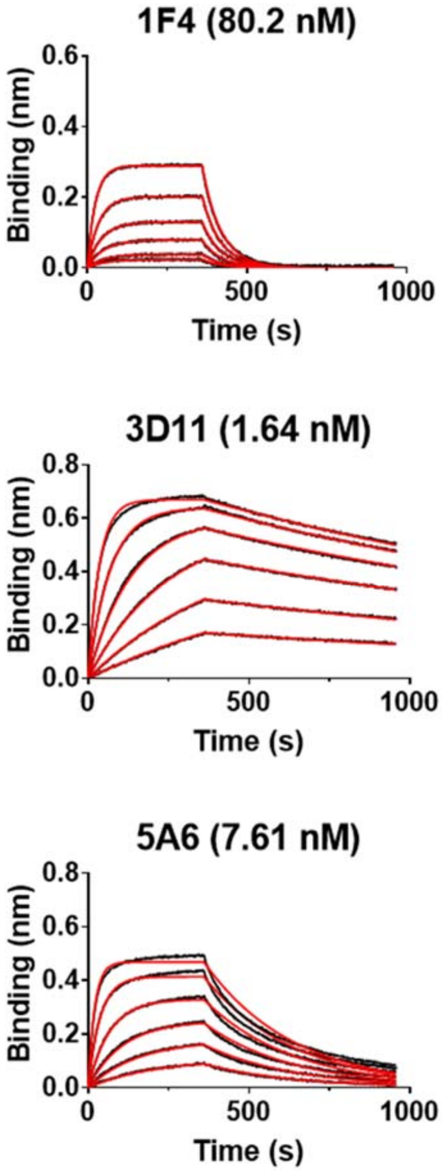

**Extended Data Fig. 3 Binding affinity of five Fab clones to SARS-CoV-2 spike RBD protein measured by biolayer interferometry.** A range of Fab concentration from 100 nM to 3.125 nM (in 2-fold dilution) was used in each experiment. The sensorgrams are in black lines and curve fittings are in red.

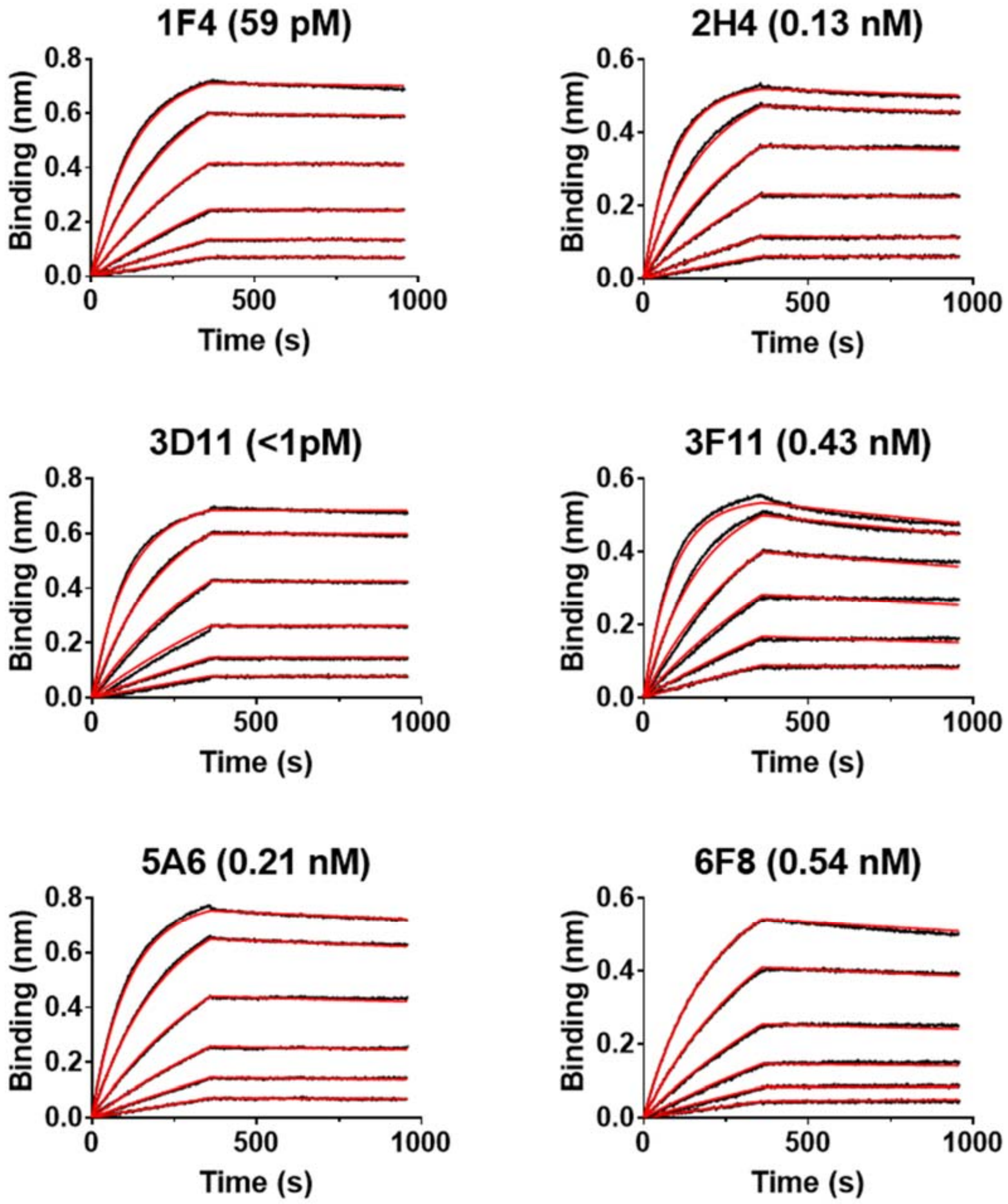

**Extended Data Fig. 4 Binding avidity of six IgGs to the RBD by biolayer interferometry.** A range of IgG concentration from 12.5 nM to 0.39 nM (in 2-fold dilutions) are shown for each IgG. The sensorgrams are in black lines and curve fittings are in red. Results shown are a representative of two independent experiments.

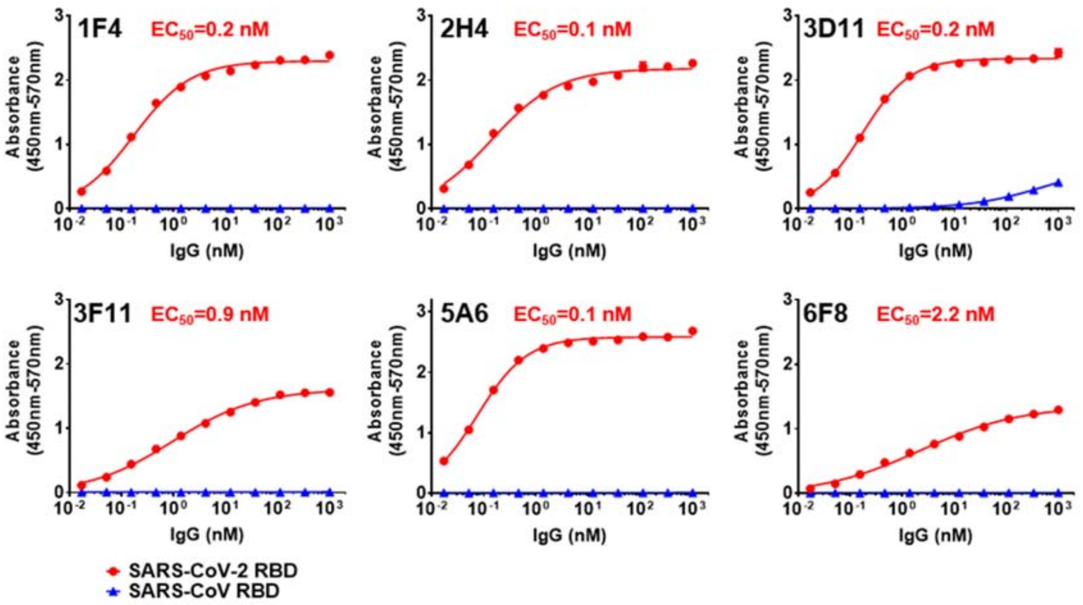

**Extended Data Fig. 5 Binding avidity of 1F4, 2H4, 3D11, 3F11, 5A6 and 6F8 IgG antibodies to SARS-CoV-2 (red circle) and SARS-CoV (blue triangle) spike RBD proteins tested by ELISA.** Data are presented as means ± SD in triplicates and are representative of two independent experiments.

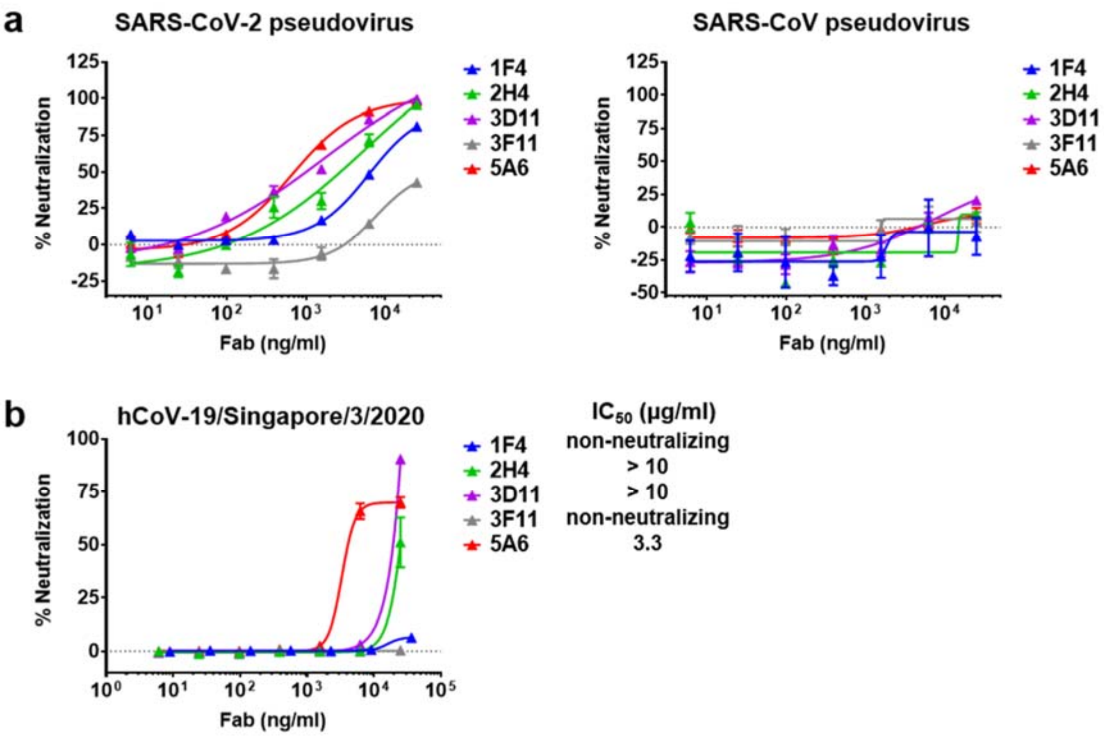

**Extended Data Fig. 6 Neutralization of the SARS-CoV-2 pseudovirus and live viruses by anti-SARS-CoV-2 spike RBD Fab antibodies. a**, Infection of CHO-ACE2 cells by SARS-CoV-2 pseudovirus (left panel) and SARS-CoV pseudovirus (right panel) were determined in the presence of 1F4, 2H4, 3D11, 3F11, and 5A6 tagless Fabs. Luciferase activities in the CHO-ACE2 cells were measured, and the percent neutralization was calculated. Data are presented as means ± SEM in triplicates and are representative of two independent experiments. **b**, Infection of Vero E6 C1008 cells by SARS-CoV-2 live virus (hCoV-19/Singapore/3/2020) were determined in the presence of 1F4, 2H4, 3D11, 3F11, and 5A6 tagless Fabs. Infection induced cytopathic effect was determined by detecting the amount of ATP present in the uninfected live cells from which the percent neutralization was calculated. Data are presented as means ± SEM in triplicates and are representative of two independent experiments.

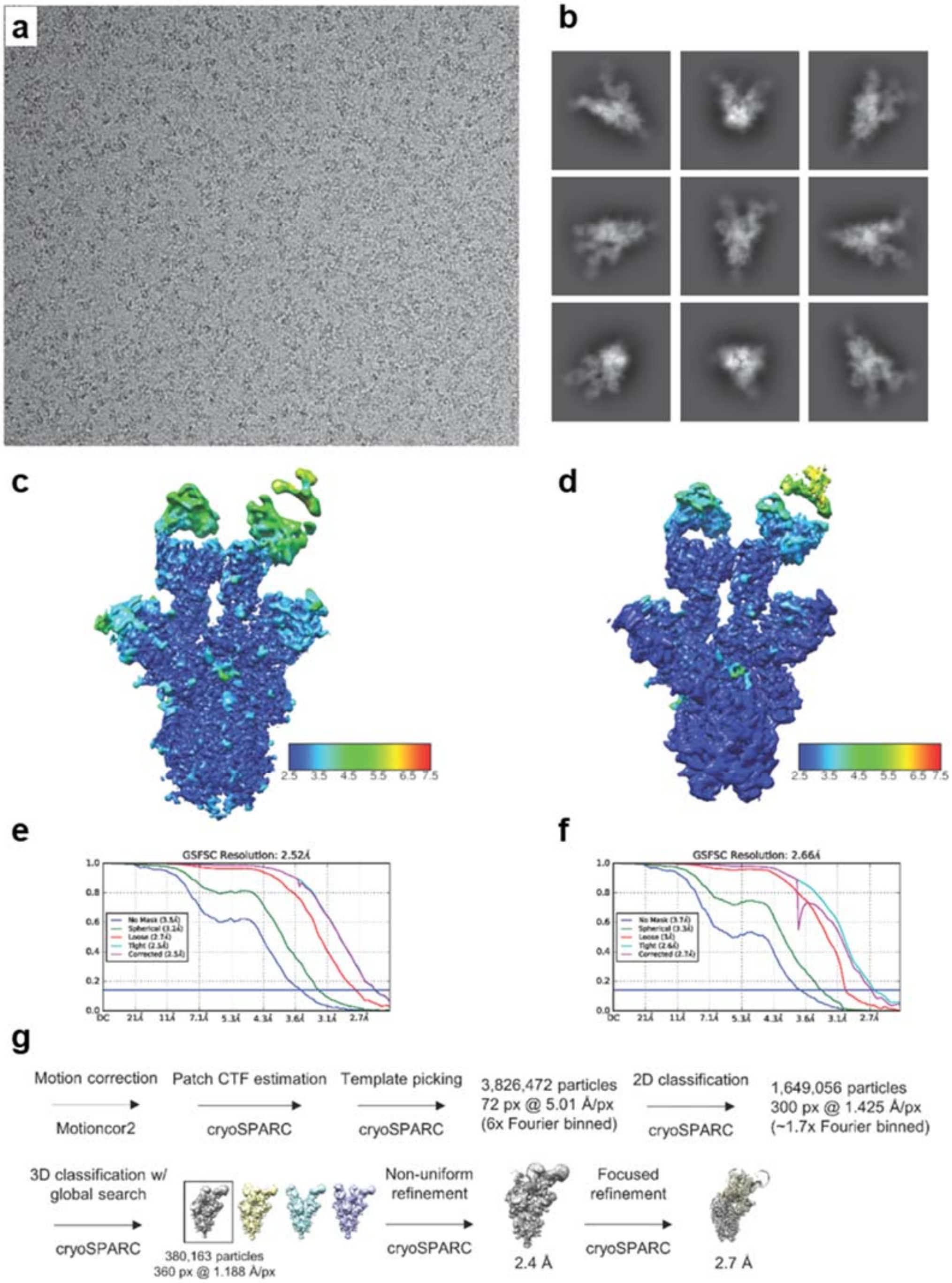

**Extended Data Fig. 7 Cryo-EM data processing. a**, Representative micrograph of spike protein-5A6 complex. **b**, Selected 2D class averages of spike protein-5A6 complex. **c**, Local resolution map of spike protein-5A6 complex from global refinement. **d**, Local resolution map of spike protein-5A6 complex from focused refinement. **e**, Gold-standard FSC curves for spike protein-5A6 global refinement. **f**, Gold-standard FSC curves for spike protein-5A6 focused refinement. **g**, Image processing pipeline employed in this work, with details for the 5A6-spike complex presented in the text.

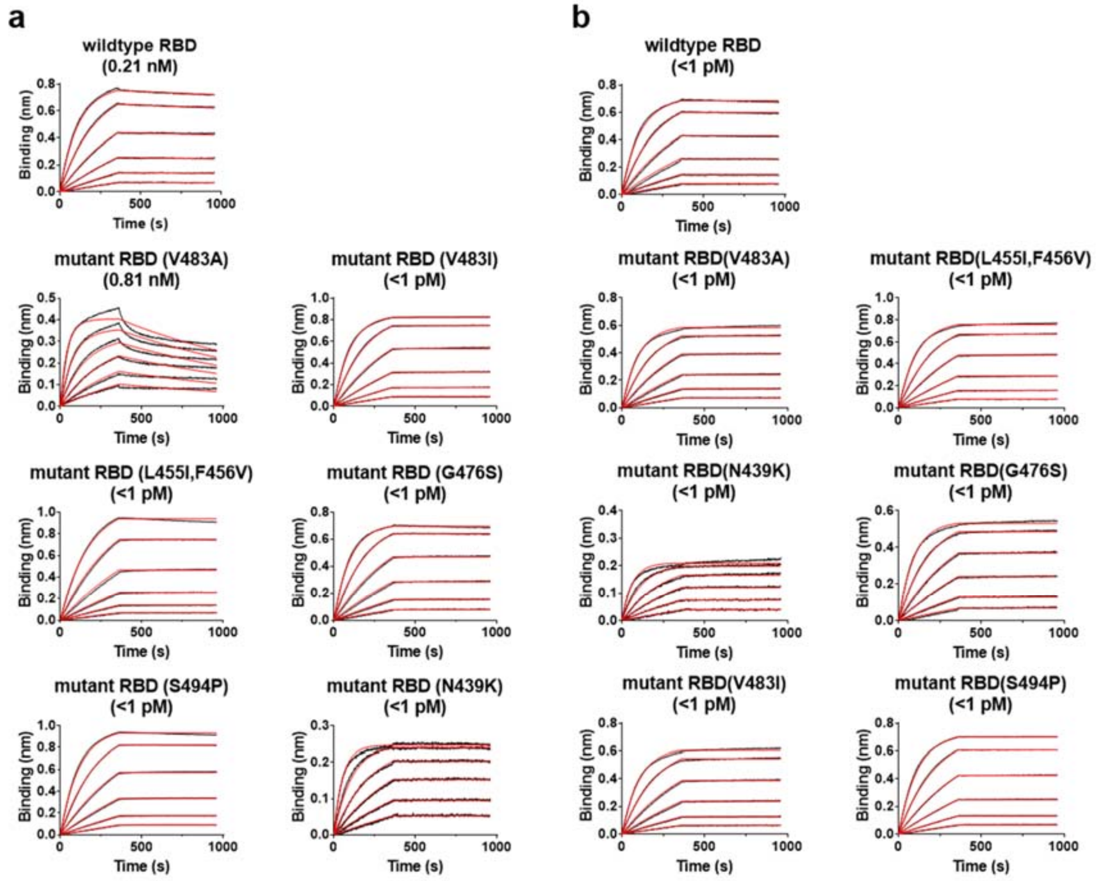

**Extended Data Fig. 8 Binding avidity of 5A6 IgG and 3D11 IgG to the wildtype RBD and RBD mutants measured by biolayer interferometry.** A range of **a**, 5A6 and **b**, 3D11 IgG concentrations from 12.5 nM to 0.39 nM (in 2-fold dilutions) are measured for each experiment. The sensorgrams are in black lines and curve fittings are in red. Results shown are a representative of two independent experiments.

**Extended Data Table 1 Cryo-EM data collection, refinement and validation statistics**

| | Spike-5A6 complex |
|---|---|
| **Data collection and processing** | |
| Magnification | EFTEM 105,000 |
| Voltage (kV) | 300 |
| Electron exposure (e–/Å$^2$) | 66.7 |
| Defocus range (μm) | -0.5-1.5 |
| Pixel size (Å) | 0.835 |
| Symmetry imposed | C1 |
| Initial particle images (no.) | 3,826,472 |
| Final particle images (no.) | 380,163 |
| Map resolution (Å) | 2.7 |
| FSC threshold 0.143 | |
| Map resolution range (Å) | |
| **Refinement** | |
| Initial model used (PDB code) | 6vyb |
| Model resolution (Å) | 3.0 |
| FSC threshold 0.5 | |
| Model resolution range (Å) | |
| Map sharpening *B* factor (Å$^2$) | -87.7 |
| Model composition | |
| Non-hydrogen atoms | 6,590 |
| Protein residues | 852 |
| Ligands | 0 |
| *B* factors (Å$^2$) | |
| Protein | 1.84/115.96/25.03 |
| Ligand | -/-/- |
| R.m.s. deviations | |
| Bond lengths (Å) | 0.012 |
| Bond angles (°) | 2.069 |
| Validation | |
| MolProbity score | 1.23 |
| Clashscore | 0.77 |
| Poor rotamers (%) | 1.22 |
| Ramachandran plot | |
| Favored (%) | 93.72 |
| Allowed (%) | 6.16 |
| Disallowed (%) | 0 |

**Supplementary Movie 1. Tour of 5A6-spike trimer 2:1 complex as shown in Figure 3.** Spike monomers are in blue (down), green (up), and red (up or down; modelled up) with 5A6 Fabs in goldenrod.
